# Medication Use is Associated with Distinct Microbial Features in Anxiety and Depression

**DOI:** 10.1101/2024.03.19.585820

**Authors:** Amanda Hazel Dilmore, Rayus Kuplicki, Daniel McDonald, Megha Kumar, Mehrbod Estaki, Nicholas Youngblut, Alexander Tyakht, Gail Ackermann, Colette Blach, Siamak MahmoudianDehkordi, Boadie W. Dunlop, Sudeepa Bhattacharyya, Salvador Guinjoan, Pooja Mandaviya, Ruth E. Ley, Rima Kaddaruh-Dauok, Martin P. Paulus, Rob Knight, Alzheimer Gut Microbiome Project Consortium

## Abstract

This study investigated the relationship between gut microbiota and neuropsychiatric disorders (NPDs), specifically anxiety disorder (ANXD) and/or major depressive disorder (MDD), as defined by DSM-IV or V criteria. The study also examined the influence of medication use, particularly antidepressants and/or anxiolytics, classified through the Anatomical Therapeutic Chemical (ATC) Classification System, on the gut microbiota. Both 16S rRNA gene amplicon sequencing and shallow shotgun sequencing were performed on DNA extracted from 666 fecal samples from the Tulsa-1000 and NeuroMAP CoBRE cohorts. The results highlight the significant influence of medication use; antidepressant use is associated with significant differences in gut microbiota beta diversity and has a larger effect size than NPD diagnosis. Next, specific microbes were associated with ANXD and MDD, highlighting their potential for non-pharmacological intervention. Finally, the study demonstrated the capability of Random Forest classifiers to predict diagnoses of NPD and medication use from microbial profiles, suggesting a promising direction for the use of gut microbiota as biomarkers for NPD. The findings suggest that future research on the gut microbiota’s role in NPD and its interactions with pharmacological treatments are needed.

## 4. Introduction

It has become increasingly clear that peripheral factors such as chemical exposome, diet, gut microbiome and lifestyle can impact brain metabolic heath and contribute to development of neuropsychiatric diseases. Several human cohorts have exhibited disruptions in the gut microbiota in neuropsychiatric disorders (NPD), including anxiety disorder (ANXD) and major depressive disorder (MDD), relative to healthy comparison (HC) subjects (reviewed in 1, 2, 3). Some of the observational differences in these cohorts have been supported by mechanistic mouse models, which raises the possibility that findings in large cohort studies can be translated into new treatments for ANXD and MDD. Proposed treatments, such as diet alteration or probiotic use, are often less expensive than current first-line treatments, and come with fewer unpleasant side effects (4). This prospect of inexpensive, non-invasive treatments for NPD has catalyzed a significant amount of research into the Gut Microbiota-Brain Axis (GMBA) over the last several years. However, there is still a significant gap between these findings and establishing a direct relationship between microbial composition and ANXD or MDD. Despite its high prevalence in both American populations and worldwide, there has been relatively little effort in studying the relationship between ANXD and the gut microbiota. Moreover, existing studies often fail to account for factors such as medication use or diet (5, 6). Although more studies have examined the relationship between MDD and the gut microbiota, these studies have similar drawbacks. Many studies have been performed outside the United States, where a larger percentage of participants are treatment-naive or have adhered to a specific treatment regimen for an extended period of time (2). Thus, these cohorts often lack the diversity of individuals living in the United States, and have significantly different pharmacological treatment courses.

Medication use, particularly selective serotonin reuptake inhibitors (SSRI) antidepressants, is relevant when discussing gut microbiota alterations in ANXD and MDD, as these medications are known to impact gut microbiota composition. However, the changes caused are debated and differ between studies. One study reported that participants using SSRI antidepressants reliably had higher abundances of *Eubacterium ramulus*, but the magnitude of this effect was small compared to differences in the microbiota seen in participants using proton pump inhibitors or metformin (7). This effect also appeared to be clustered in individuals using paroxetine, suggesting it could be specific to that particular SSRI. Another study reported a decrease in *Turicibacteraceae* abundance in SSRI-users, which was also supported by a mouse study in which mice were given fluoxetine (8, 9). The observation that fluoxetine decreases the relative abundance of *Turicibacter sanguinis* is particularly interesting because *T. sanguinis* imports serotonin (5-HT) through an analog of the 5-HT transporter (SERT), which fluoxetine inhibits (9). Therefore, these gut microbiota changes could also influence metabolic pathways.

SSRI use is known to modulate levels of acylcarnitines, amines, lipids, and bile acids (BA; 10, 11). Specifically, levels of the primary BA chenodeoxycholic acid (CDCA) were lower in severely depressed and anxious participants, while levels of secondary BAs produced from CDCA, such as lithocholic acid were significantly higher in these individuals (11). The high level of bacterially produced secondary bile acids derived from LCA and metabolism of CDCA correlated with anxiety severity. Given that members of the microbiota are known to modulate the bile acid pool, including making previously unknown conjugated bile acids, it is possible that shifts in the gut microbiota could be behind these shifts in the metabolome in ANXD and MDD (12, 13). In a similar fashion, treatment with citalopram/escitalopram reduced serotonin levels and increased concentrations of indoles, which are known to be associated with microbial cometabolism (14). While another study notes that levels of the gut microbiome-derived metabolite indoxyl sulfate are elevated in participants with depression both at baseline and after successful treatment with medications, this does rule out the possibility that other microbially-modified indoles are altered in MDD or ANXD (15). The rapid acting antidepressants ketamine and esketamine modify levels of indole-3-acetate and methionine, further implicating tryptophan metabolism in MDD, and highlighting the importance of microbes that modify tryptophan (16).

While some literature reports that SSRIs return the microbiota to a “normal state” after depression alters the gut microbiota composition, these conclusions are supported by beta diversity significance statistics rather than differential abundance of specific taxa (17, 18). For example, one study reports that because a subgroup of participants had microbiota that resembled those of HC after taking escitalopram, their microbiota was returning to normal, even though their predicted functional profiles were significantly different (17). Another study similarly reported that since there was no difference in beta diversity between HC and participants with MDD after eight weeks of vortioxetine treatment, and a subset of the taxa enriched in MDD were lowered after SSRI treatment, that the medication is acting through alternation of gut microbiota composition (18). While possible that SSRIs modify some taxa of the gut microbiota specifically, particularly those that express SERTs, the conclusion that the entire microbiota trends towards normal is not supported by the relative abundance data reported. Other studies support this conclusion as well, such as one in which Prevotella and Klebsiella are found to be specific markers of MDD in participants taking SSRIs compared to controls (19). Even though some specific taxa may be exclusive to a certain locale, differences in specific taxa are generally more robust than differences reported at a summary level (i.e. beta diversity) in other neuropsychiatric diseases (20). This suggests that differential abundance analyses are preferable.

Aside from alterations specific to medications, anxiety and depression have also independently been associated with changes in the gut microbiota. One study, for instance, examined unmedicated participants with either ANXD or MDD and identified specific OTUs that differentiated ANXD from MDD (21). This is significant given that the disorders have several common phenotypes and can be difficult to distinguish in a clinical setting. Another study was able to discriminate between participants with MDD and HC using either taxonomic or functional labels (22). In general, most studies of microbiota differences between ANXD and/or MDD and HC report reductions in the relative abundances of anti-inflammatory bacteria in individuals with diagnosed ANXD and/or MDD (23, 24).

While the aforementioned studies mentioned are informative, many have very small sample size (<50 participants), limiting their statistical power. Furthermore, few have been performed on an American population with a wide range of medication types and dosages, limiting their generalizability. Here, we sought to identify gut microbiota differences specific to ANXD and MDD, as well as those specific to anxiolytic and antidepressant use, in a large United States-based cohort with high utilization of medications. We hypothesized that ANXD and MDD would each be associated with distinct microbiota compositions. Further, we predicted that the medications used to treat these conditions would also be associated with particular microbial features. These hypotheses would have significant implications for both the use of gut microbiota as a biomarker of affective disorders and a supplement or alternative to pharmacological interventions.

## 5. Materials / Subjects & Methods

### Description of cohort & how filtering was performed

Stool samples collected from participants in the Tulsa-1000 and NeuroMAP CoBRE cohorts were analyzed. Full details describing the Tulsa-1000 study can be found in Victor et al and a description of the NeuroMAP CoBRE study is in Kuplicki et al (25, 26). Participants were given stool sample collection kits (BD SWUBE Dual Swab Collection System), which were returned to a study coordinator at the Laureate Institute for Brain Research (LIBR) within one day of collection. Samples were stored at −80 °C, until DNA extraction was performed using the MO BIO PowerMag Soil DNA Isolation Kit and the Earth Microbiome Project’s (EMP) protocols (27). Amplicons of the V4 region of the 16S rRNA gene were generated using barcoded EMP primers as described here (28). Paired-end 150bp reads were generated by sequencing the amplicons on an Illumina MiSeq at the Knight Lab (29). A subset of DNA extracts was sequenced using shotgun metagenomic sequencing at the Max Planck Institute for Biology Tübingen. Library preparation was performed using a Tn5 purification and tagmentation protocol, and paired-end 150bp reads were sequenced on an Illumina HiSeq 3000 (30).

Study staff at LIBR administered self-report assessments to the individuals to collect information on demographic, clinical, and psychiatric features. The full set of assessments can be found in Victor et al (25). Our study relied on the evaluation of recent medication use, categorized by their Anatomical Therapeutic Chemical (ATC) codes, ANXD and/or MDD diagnosis via the MINI International Neuropsychiatric Interviews version 6 or 7, and severity of ANXD and/or MDD symptoms with the Overall Anxiety Severity and Impairment Scale (OASIS) score and Patient Health Questionnaire-9, respectively (31–34). Ethical approval for the both studies was obtained from Western Institutional Review Board. All methods were carried out in accordance with relevant guidelines and regulations. Informed consent was obtained by trained members of the research team.

Raw data were uploaded to Qiita [DOI: 10.1038/s41592-018-0141-9] under study ID 10424 (35). Adapters were removed, then the data were trimmed to 150bp using fastp (36). The resulting 16S reads were processed using Deblur version 2021.09, retaining the positively-filtered feature tables (37). In addition, only amplicon sequencing variants (ASVs) that mapped to the Greengenes2 2022.10 database were retained (38). Human reads were filtered out from the metagenomics data with Minimap2 using the reference genomes: GRCh38, CHM13-T2Tv2.0, and Human Pangenome (39, 40). Data were aligned to the Web of Life v2 reference database using Bowtie2 (version 2.4.2) with parameters adapted from SHOGUN (41–43). Aligned sequences were processed into genomic operational taxonomic units (gOTUs) using Woltka (version 0.1.4) (44).

Filtered 16S feature tables from the following Qiita preps were merged: 1428, 2145, 2569, 4004, 4607, 4685, 6173, 7243, and 8287. The resulting merged table had 1440 fecal samples with matching metadata, and 1008 of these fecal samples matched individuals with LIBR metadata entries. However, some participants donated multiple samples; to choose the “best-sequenced” sample, the remaining fecal samples were sorted by their sampling depth and the sample that had the highest read depth per individual was retained (leaving 738 samples). Only samples that had over 5000 counts were kept (704 samples retained). The data were then filtered so that only participants diagnosed with ANXD or MDD and HC remained (700 samples). Participants who used antibacterial medications were also removed from the sample set (666 samples retained).

Filtered metagenomic preps from the following Qiita preps were merged: 16292, 16293, 16297, 16298, 16299. The resulting merged table had 321 fecal samples, of which 296 had matching LIBR metadata. Only samples that had over 500,000 reads were kept, retaining 271 samples.

### Determination of effect size distributions

Alpha and beta diversity metrics were calculated using QIIME 2’s diversity and gemelli plugins (version 2022.11) (45, 46). The Greengenes2 2022.10 tree was used for phylogenetic metrics (38). ASV feature tables were rarefied to 5000 sequences per sample over ten iterations, then distance matrices were constructed. Independent categorical effect sizes were calculated using scikit-bio’s implementation of PERMANOVA (47). We were interested in exploring the interaction between diagnostic status and medication use, given that medications are almost exclusively used by participants with at least one NPD diagnosis. Therefore, the cohort was divided into medicated and unmedicated participants for effect size calculations. The effect of multiple variables on microbial beta diversity was calculated using vegan’s adonis function (48). The order of variables was determined by their independent effect size, since adonis proceeds in order.

### Identifying differential ASVs & subsequent log ratio calculation

Differential taxa were identified using Bayesian Inferential Regression with BIRDMAn (49). Given that medication use may alter gut microbiota composition, only unmedicated participants were included in the differential abundance analyses. Age, sex, cohort (CoBRE or Tulsa-1000), and BMI were included as covariates. Features were considered credibly associated with a phenotype if their Bayesian credible interval contained only negative values or only positive values. The former was assigned as “negatively associated” with the phenotype and the latter was assigned as “positively associated.” A log ratio was then constructed for the phenotype as the natural logarithm of the sum of all “positively associated” taxa divided by the sum of all “negatively associated” taxa. If the sum of either positively or negatively associated taxa was zero, the sample was not assigned a ratio for that phenotype.

### Transfer of ASV-level results to American Gut Project data

To determine whether the results found in the Tulsa cohort could apply to another large American cohort, we attempted to validate our results on participants of similar ages in the American Gut (AGP) cohort (50). To apply the top differential ASVs to the AGP data, all samples for individuals aged fifty-five and below were retrieved using the redbiom query “where `(qiita_study_id == 10317) AND (host_age <= 55)”` in the “`Deblur_2021.09-Illumina-16S-V4-150nt-ac8c0b”` context (51). Samples were subsequently filtered using the following criteria: individuals below the age of eighteen were removed, non-stool samples were removed, ASVs associated with blooms previously observed in the American Gut Project were removed, for subjects with multiple samples a single arbitrary one was retained, and samples with fewer than 5000 counts were removed (52). The MDD log ratio was then generated using the credible taxa from the Tulsa data. Kruskal-Wallis tests were performed to assess significance.

### Transfer of ASV-level results to shallow shotgun metagenomics data

Given our limited sample size in the shotgun metagenomic samples, we attempted to repurpose our BIRDMAn differentials in the whole-genome sequencing (WGS) data. To do so, a custom Python script was used to identify the nearest node between each ASV and gOTU on the Greengenes2 phylogenetic tree (38). Dictionaries were then created to map between ASVs and gOTUs; these were utilized to “transfer” the differential ASV results to the shotgun metagenomics data.

### Random Forest Classification

Random forest classifiers were constructed with scikit-learn version 0.25.3 to predict NPD diagnosis and medication use from microbial features (53). Stratified k-fold cross-validation was used to assign five training and testing sets for each classifier. Each classifier was run ten times to generate multiple predictions. Additionally, four feature selection schemes were employed: subsetting to microbes with 5% or higher prevalence, microbes identified through ridge regression, microbes with positive feature importance scores (across any fold, as feature selection was performed separately for each fold), and microbes identified as significantly associated with NPD diagnosis or medication use through BIRDMAn.

### Data & Code Availability

16S and Metagenomic sequencing data are available under study ID 10424 in Qiita (35). Scripts used for analyses in this manuscript can be found at: https://github.com/ahdilmore/Tulsa1000_microbiome.

## 6. Results

### Cohort Characteristics

After performing quality control filtering (removing duplicate individuals, removing samples with low read count, matching to metadata), 666 fecal samples remained. These matched 164 HC and 502 individuals with either an ANXD or MDD diagnosis. A majority of participants were diagnosed with both ANXD and MDD (n=301; 60.0%). Both the ANXD and MDD groups were composed of more females (77.71% and 74.52%, respectively) compared to the healthy control group (63.41%). This cohort reported widespread medication use, including but not limited to, antidepressants (ATC N06A; 30.63%) and anxiolytics (ATC N05B; 11.86%). Full cohort information can be found in **Table 1**.

**Table 1.** Cohort Characteristics: Demographic information of the subset of the Tulsa cohort with matching fecal samples that passed quality control (n=666).

|  | **ANXD** | **MDD** | **HC** |
| --- | --- | --- | --- |
| **Number** | 332 | 471 | 164 |
| **% Diagnosed with both**<br>**ANXD and MDD** | 90.66 | 63.91 | N/A |
| **Age** | 33.41 ± 10.52 | 34.17 ± 11.02 | 30.3 ± 10.72 |
| **Body Mass Index** | 28.04 ± 5.48 | 28.39 ± 5.55 | 26.71 ± 5.25 |
| **Sex (% Female)** | 77.71 | 74.52 | 63.41 |
| **% Using Anxiolytics**<br>**(ATC: N05B)** | 18.98 | 14.23 | 0 |
| **% Using Antidepressants**<br>**(ATC: N06A)** | 40.36 | 40.76 | 1.22 |

### Microbial Diversity Analysis

After accounting for multiple comparisons, there were no significant differences in alpha diversity between ANXD and MDD diagnosis groups, between males and females, or between anxiolytics and antidepressants use (**[Table S1](https://drive.google.com/file/d/1s0JBGsIKDMRmkiXya7Rr4jSkBKcSHQdi/view?usp=sharing)**). While MDD diagnosis and antidepressant use, both were significantly associated with beta diversity, antidepressant use exhibited a larger effect size on beta diversity overall (**Fig 1A-B**). In addition, anxiolytic use, sex, and cohort were significantly associated with beta diversity (**Fig 1C**). In unmedicated participants, only sex remained significantly associated with beta diversity and in medicated participants, only cohort and sex remained significantly associated with beta diversity (**Fig 1D-E**). When we utilized the adonis function, incorporating medication use and NPD diagnosis in the same formula, we found that antidepressant use exhibited a significant effect, while ANXD and MDD diagnosis did not (**Fig 1F**). Full beta diversity statistics are available in **[Table S2](https://drive.google.com/file/d/1vvwkyJU6QCv_kjnvii1U_x3Ohl82bta6/view?usp=sharing)**.

**Figure 1.**
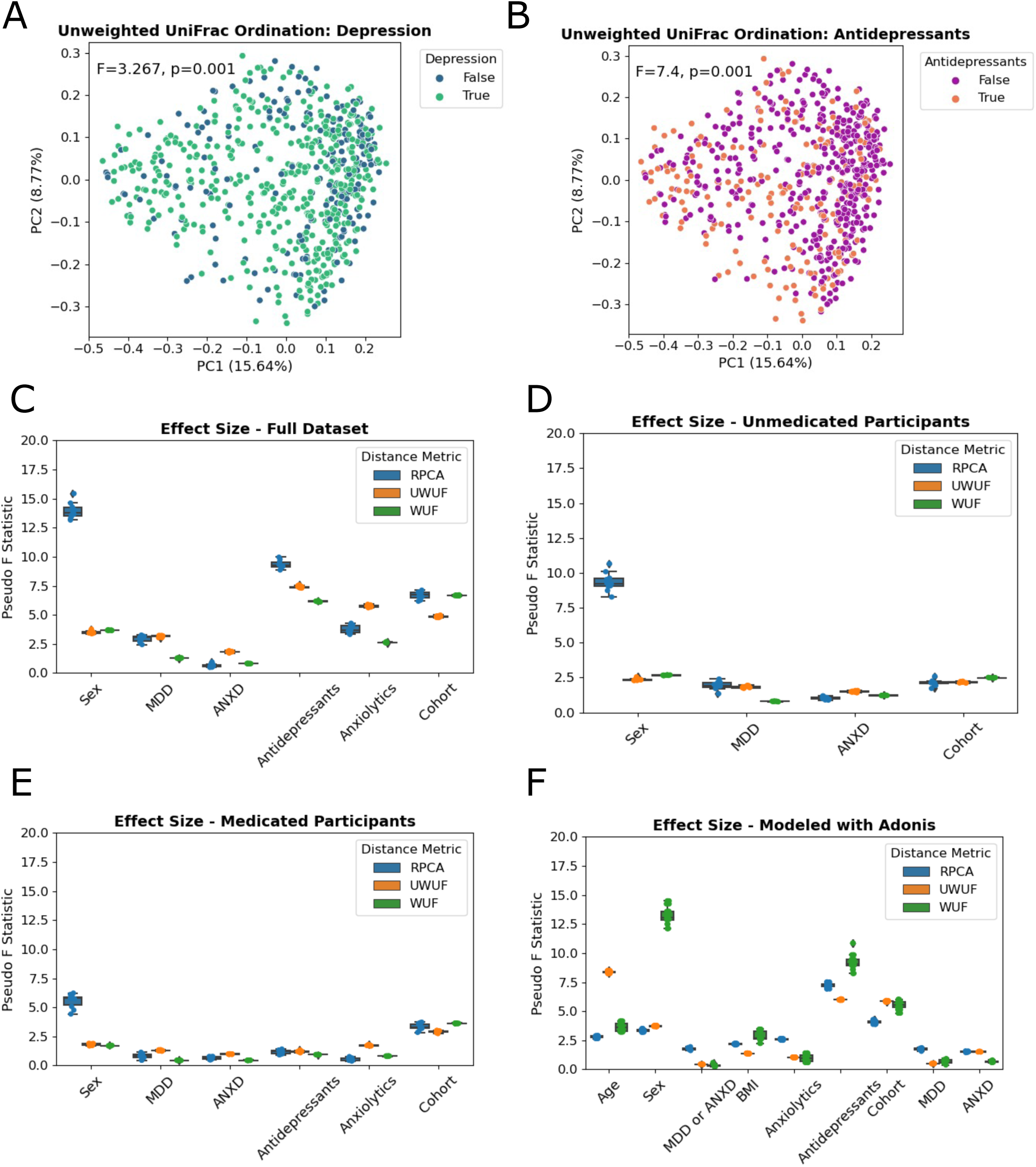
Medications Have a Stronger Effect on Gut Microbiota Beta Diversity than ANXD or MDD Diagnosis. **(A)** Principal coordinates (PCoA) plot of Unweighted UniFrac distances shows statistically significant, but minimal visible separation between participants with MDD and HC. **(B)** The same plot colored by antidepressant use shows a stronger statistical and visual separation between participants using antidepressants and those not using them. **(C)** Independent effect sizes of (categorical) demographic and psychiatric variables on gut microbiota beta diversity across the entire cohort. **(D)** Independent effect sizes of (categorical) demographic and psychiatric variables on gut microbiota beta diversity across only unmedicated individuals. **(E)** Independent effect sizes of (categorical) demographic and psychiatric variables on gut microbiota beta diversity across only medicated individuals. **(F)** Effect sizes of categorical and quantitative variables were modeled together using the following formulas in vegan’s Adonis function: Antidepressant Use + Cohort + Anxiolytic Use + Sex + Age + BMI + MDD or ANXD / MDD / ANXD.

### Identification of Microbes Associated with ANXD, MDD, and Medication Usage

#### Microbes associated with ANXD

When we compared 172 unmedicated participants with ANXD diagnoses to 152 unmedicated HC (**Fig 2A**), we found 94 ASVs, of which 44 were significantly associated with ANXD and 50 ASVs significantly associated with HCs (**Fig S1A-B**). For illustrative purposes, the top 15 ASVs associated with ANXD and HC are shown in **Fig 2B**. We computed log ratios of the top microbes associated with ANXD in unmedicated participants, and observed higher ratios in both participants with ANXD alone and those with MDD alone, as compared to HC (**Fig 2C**). These ratios were also significantly higher in participants with both ANXD and MDD, as compared to participants with only MDD. However, there was no significant difference between individuals with ANXD and those with MDD. We also found that the ANXD-associated log ratio was significantly lower in medicated participants with ANXD (**Fig 2D**). This log ratio was also correlated with anxiety severity, as measured by the OASIS score (**Fig 2E**).

**Figure 2.**
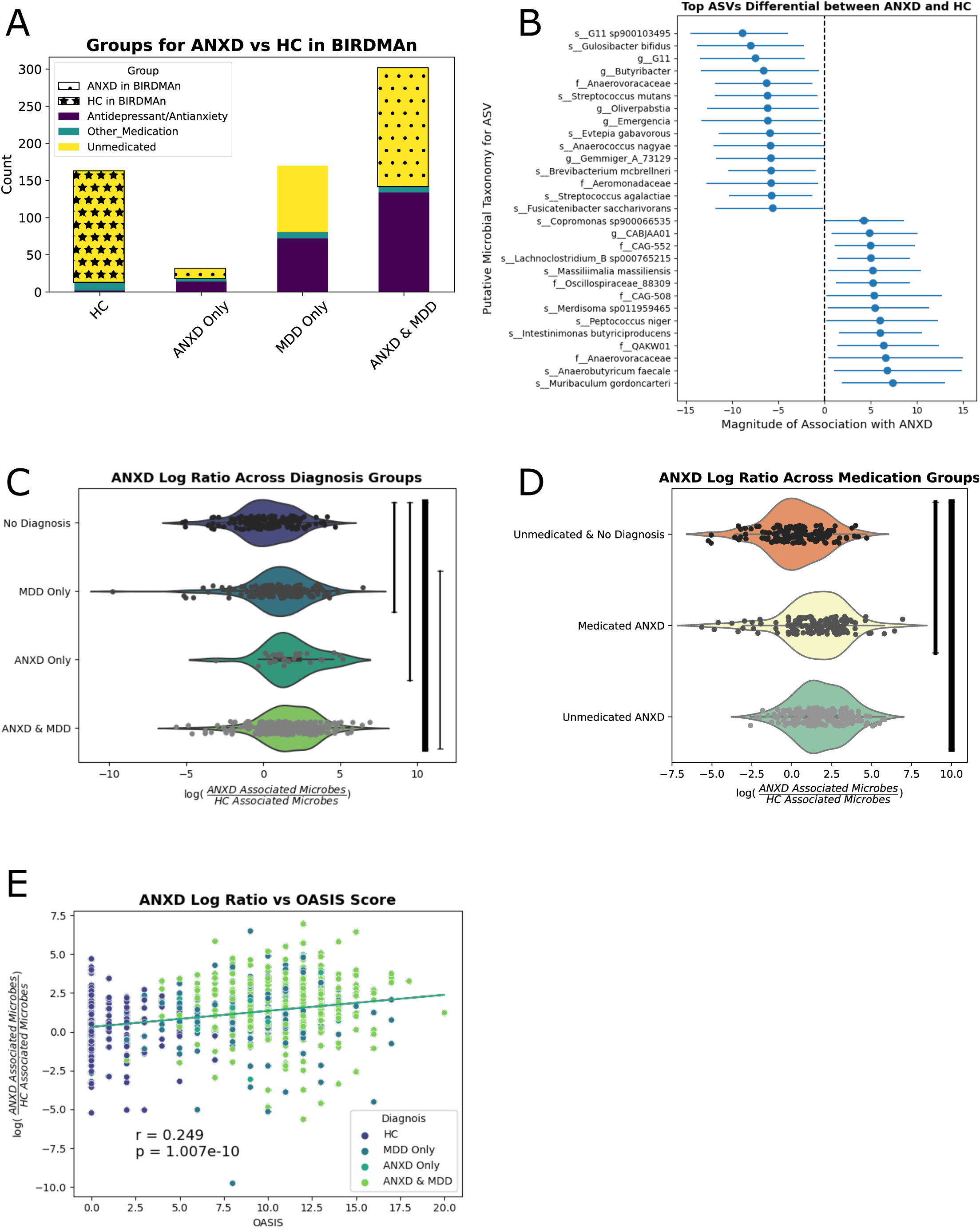
Identification & validation of microbes enriched in ANXD: **(A)** Visualization of the participants used in the discovery cohort. Unmedicated participants with ANXD were compared to unmedicated HC. **(B)** The top 15 microbes credibly associated with ANXD or HC are displayed. **(C)** Log ratios were created comparing the relative abundance of all credible microbes associated with ANXD to all credible microbes associated with HC. Thickness of the line is correlated with the negative log of the p-value between the two groups. **(D)** Log ratio displayed by medication category. **(E)** Scatterplot between ANXD log ratio and OASIS score.

#### Microbes associated with MDD

When we compared 248 unmedicated participants with a MDD diagnosis to 152 unmedicated HC (**Fig 3A**), we found 47 microbes that were significantly associated with MDD and 64 microbes significantly associated with HC (**Fig S1A-B**). The top 15 associated with either group are shown in **Fig 3B**. We constructed log ratios of the top microbes associated with MDD in unmedicated participants, and found that this ratio was higher in participants with any psychiatric diagnosis compared to HC (**Fig 3C**). However, this log ratio was not significantly different between MDD and ANXD or between participants with both diagnoses and participants with only a MDD diagnosis. The MDD log ratio was higher in both medicated- and unmedicated participants (**Fig 3D**). The MDD specific log ratio was significantly correlated with depression severity, as measured by the PHQ-9 questionnaire (**Fig 3E**). There was a significant difference in the log ratio between participants who self-reported “Diagnosed by a medical professional (doctor, physician assistant)” than participants who self-reported “I do not have this condition” in the American Gut cohort (**Fig 3F**).

**Figure 3.**
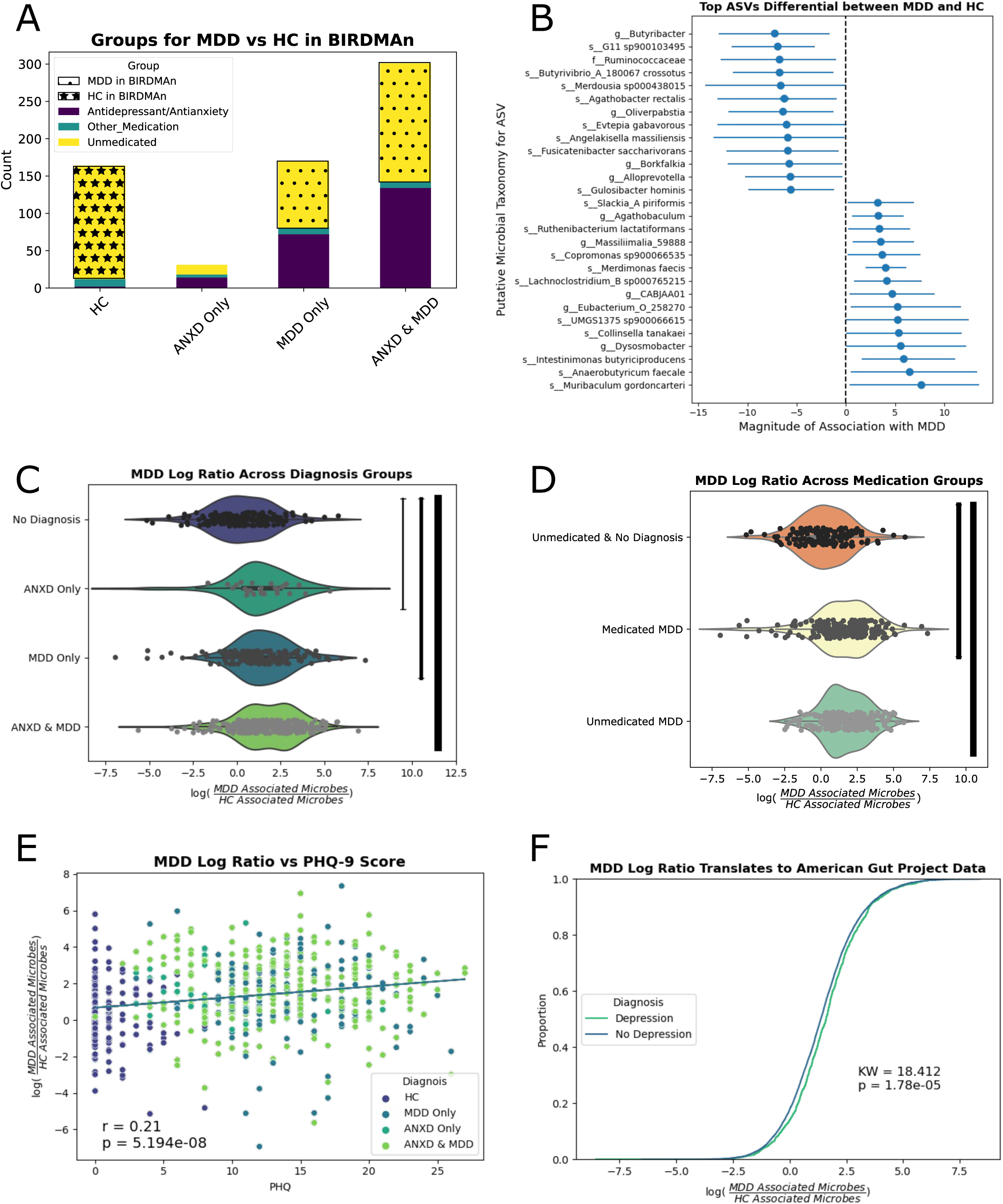
Identification & validation of microbes enriched in MDD: **(A)** Visualization of the participants used in the discovery cohort. Unmedicated participants with MDD were compared to unmedicated HC. **(B)** The top 15 microbes credibly associated with MDD or HC are displayed. Log ratios were created comparing the relative abundance of all credible microbes associated with MDD to all credible microbes associated with HC. **(D)** The same log ratio displayed by medication category. **(E)** Scatterplot between MDD log ratio and PHQ-9 score. **(F)** Empirical cumulative distribution plot between the MDD log ratio in American Gut participants with or without a self-reported MDD diagnosis.

#### Microbes associated with medication use

When we compared 172 unmedicated participants with an ANXD diagnosis to 79 participants taking anxiolytics and an ANXD or MDD diagnosis (**Fig 4A**), we found 77 microbes that were significantly associated with no medication use and 88 microbes significantly associated with anxiolytic use (**Fig 1C-D**). The top 15 microbes associated with either group are shown in **Fig 4B** for illustrative purposes. When we compared 248 unmedicated participants with a MDD diagnosis to 204 participants taking antidepressants and a MDD or ANXD diagnosis (**Fig 4D**), we found 73 microbes that were significantly associated with unmedicated participants with MDD and 102 microbes significantly associated with antidepressant use (**Fig S1C-D**). The top 15 associated in either direction are shown in **Fig 4C**. After constructing log ratios for anxiolytic and antidepressant use, we found that the anxiolytic log ratio was significantly enriched in participants using antidepressants, anxiolytics, and both types of medications. The difference between anxiolytics and antidepressants, as well as the difference between anxiolytics and antidepressants and antidepressants alone was significant (**Fig 4E-F**).

**Figure 4.**
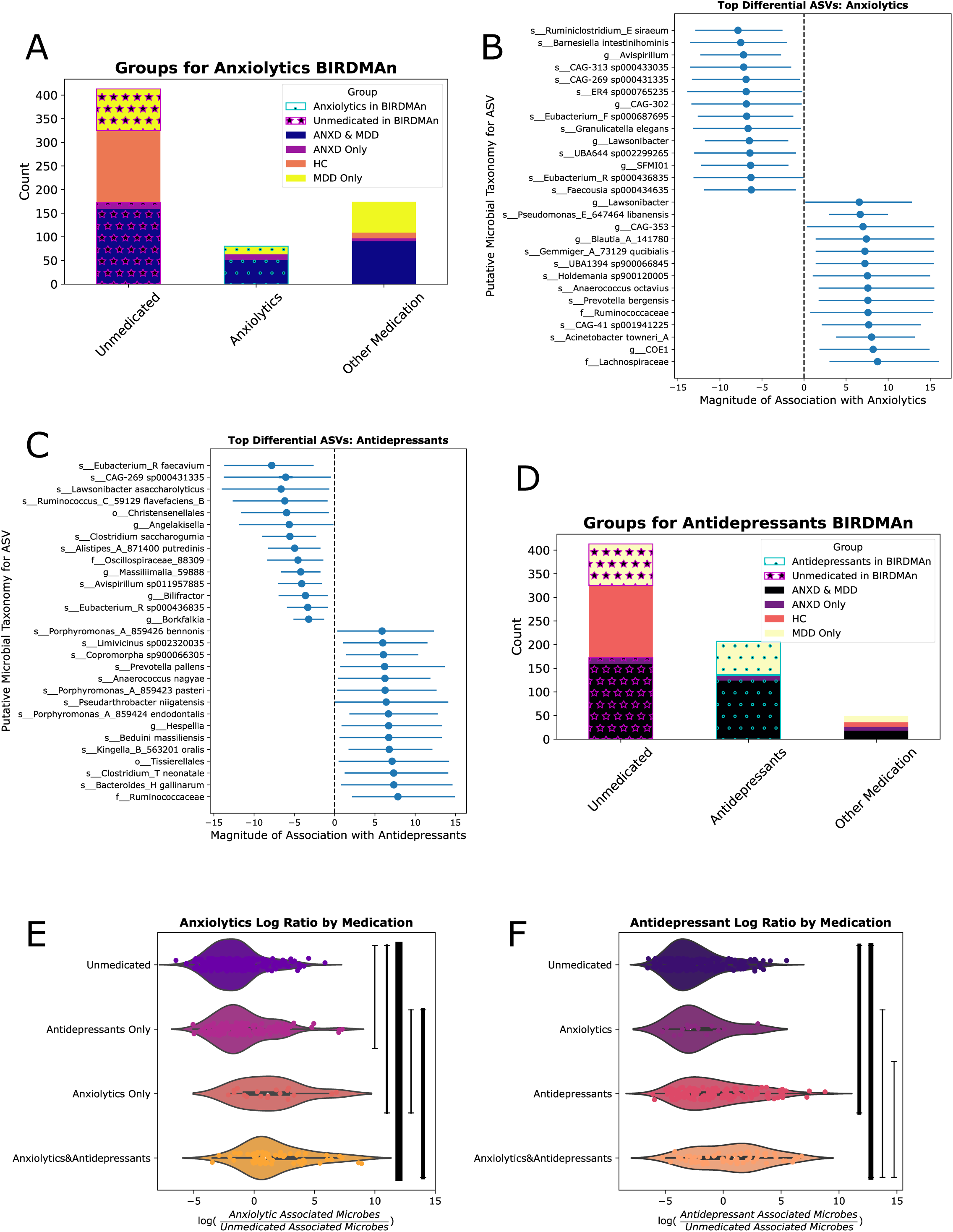
Identification & validation of microbes enriched with anxiolytic and/or antidepressant use: **(A)** Visualization of the participants used in the discovery cohort. Unmedicated participants with an ANXD or MDD diagnosis were compared to individuals with an ANXD or MDD diagnosis who were taking anxiolytics. **(B)** The top 15 microbes credibly associated with anxiolytics or no medication use are displayed. **(C)** Visualization of the participants used in the discovery cohort. Unmedicated participants with an ANXD or MDD diagnosis were compared to individuals with an ANXD or MDD diagnosis who were taking antidepressants. **(D)** The top 15 microbes credibly associated with antidepressants or no medication use are displayed. **(E)** Log ratios were created comparing the relative abundance of all credible microbes associated with anxiolytics to all credible microbes associated with no medication usage. **(F)** Log ratios were created comparing the relative abundance of all credible microbes associated with antidepressants to all credible microbes associated with no medication usage.

All comparison statistics are available in **[Table S3](https://drive.google.com/file/d/1FmmcuBonMvf9Ae2OCJ98YBHFLDOOvvGX/view?usp=drive_link)**.

### Classification of Psychiatric Diagnosis and Medication Usage from Microbial Profiles

We reliably classified participants with ANXD, MDD, and those with both ANXD and MDD from HC using microbial taxa alone. This held true for both the entire group of participants and when only unmedicated participants were considered (**Fig 5A-B**). In all instances, feature selection using microbes significantly enriched by BIRDMAn markedly enhanced classification accuracy (measured by the area under the receiver operating characteristic curve (AUC)) compared to other feature selection methods (**Fig 5A-B**). In both the entire cohort and among unmedicated participants alone, the average precision score (APR) consistently surpassed the population’s true positive rate, indicating that the classification was both accurate and precise (**Fig 5C-D**).

**Figure 5.**
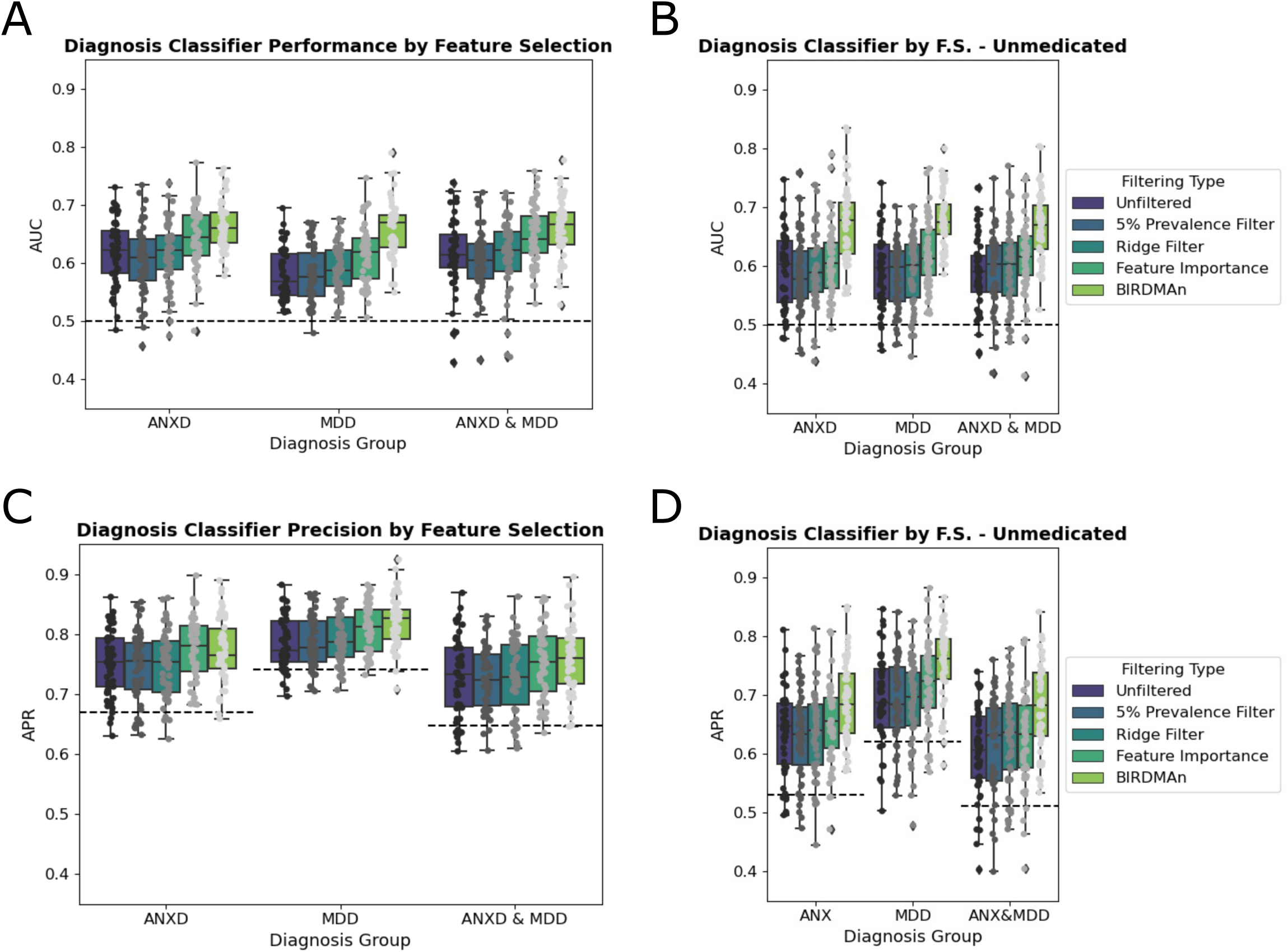
Random Forest Classifiers demonstrate high accuracy and precision when predicting ANXD and/or MDD diagnosis. **(A)** Area under the receiver operating characteristic curve (AUC) is displayed for a diagnosis classifier distinguishing ANXD diagnosis, MDD diagnosis, or ANXD and MDD diagnosis from HC. Four types of feature selection were applied in addition to the raw data. **(B)** Random Forest Classifier was applied to only unmedicated participants. **(C)** Average precision (APR) scores were also calculated across the different classifications and feature selection methods. **(D)** APR scores in the unmedicated subset.

The diagnosis classifier also consistently yielded AUCs above 50% in the AGP cohort, and APRs consistently exceeded the true positive rate in the population (**Fig S4A-B**). This suggests that the classification was both precise and accurate, even considering the imbalanced nature of the AGP dataset.

The medication classifier performed significantly better than chance for anxiolytics, antidepressants, and both, but precision scores were lower than the true positive rate in the population (**Fig S4C-D**). BIRDMAn-selected features only significantly improved AUCs for anxiolytics, whereas these selected features improved AUCs across the board for the diagnosis classifier (**Fig S4C**).

## 7. Discussion

This study aimed to elucidate the relationship between the gut microbiota, NPD, and medication use. Through rigorous quality control measures, 666 fecal samples from the Tulsa cohort were analyzed: 164 healthy comparison subjects and 502 individuals diagnosed with ANXD, MDD, or both. There were five main results. First, there was no significant difference in alpha diversity across NPD groups, sex, or medication use. Second, antidepressant use emerged as a major influence on gut microbiota beta diversity, surpassing the effect size of MDD diagnosis. Third, several microbes associated with ANXD and MDD independent of medication use were identified through Bayesian Inferential Regression. Fourth, participants taking antidepressants had a higher abundance of a small subset, but not the majority, of microbes that are typically depleted in MDD, highlighting a potential interaction between pharmacological treatments and gut microbiota. Finally, machine learning classifiers were able to label NPD diagnosis, but not medication use, accurately and precisely from microbial profiles, suggesting that it is feasible to use gut microbiota as biomarkers for NPD. Taken together, the study highlights the significant influence of medication use on gut microbial diversity and identifies specific microbes associated with ANXD and MDD, suggesting avenues for non-pharmacological interventions. The findings advocate for further research to explore the role of gut microbiota in NPD and its interaction with pharmacological treatments.

These results support the findings of several other studies that there is a unique microbiota signature associated with ANXD and MDD. For example, the Texas Resilience Against Depression identified a network of microbes that commonly co-occur in participants with MDD (54). These signatures are not exclusive to the stool microbiota: another study identified negative associations between the *Bacteroidaceae* family and *Bacteroides* genus with depression symptoms, along with positive associations between the *Desulfovibrionaceae* family and *Clostridiales* Family XIII with anxiety symptoms in the serum microbiota, while another found links between members of the oral microbiota and MDD (55, 56). These studies suggest that the GMBA may extend beyond microbes in the gut, indicating the potential existence of unique microbial signatures of ANXD and MDD throughout the body. Consequently, potential biomarkers for these conditions may be available at more easily accessible sites.

Other studies have also examined the interactions between the GMBA and other physiological conditions. For example, *Clostridium* was found to be higher in individuals with comorbid generalized anxiety disorder and functional gastrointestinal disease, suggesting that some microbial perturbations could be influenced by changes from other diseases (57). Similarly, in a study investigating sleep quality, gut microbiota, and metabolome profiles in patients with ANXD and MDD, *Bacteroides* was found to be positively correlated with sleep quality scores. This suggests that particular microbes are depleted in participants with insomnia relative to people with ANXD and MDD but not insomnia (58). Studies like these support our examination of particular microbes associated with medication use because, while some microbes may be associated with ANXD or MDD overall, there is also a subset of microbes associated with particular phenotypes. The discovery of these associations between microbes and sub-phenotypes may aid in developing precision treatments.

In terms of medication use, one study found that psychotropic medication use decreases the alpha diversity of the gut microbiota in patients with ANXD and MDD, particularly with higher medication doses (59). While our work does not support this particular finding, this study does support the hypothesis that psychotropic medications alter the microbiota in NPD. This description of microbiota diversity underscores the complexity of the relationship between psychiatric medications and the microbiota, as it does not consider specific taxa. However, work like this does raise the question of whether medication-induced alterations in the microbiota could influence treatment outcomes for NPD. One study addresses the potential for probiotic therapy to augment ANXD and MDD treatments, by supplementing with *Lactobacillus plantarum JYLP-326* (60). While this addresses the ability of microbial treatment to manage symptoms of ANXD and MDD, it does not discuss how probiotics may be used in conjunction with other traditional treatments for the conditions, especially since these medications may further alter the microbiota themselves.

However, the sources of GMBA disruption in NPD are poorly characterized. Although several sources report disturbances to the GMBA in ANXD and MDD it is unclear whether these changes are the result of the condition itself, the medications taken for the conditions, or other factors such as diet. Our study tackles this question by identifying in a large cohort of American individuals, specific microbes that are associated with ANXD, MDD, anxiolytic use, and antidepressant use. We used a Bayesian Differential Ranking algorithm that allows us to account for the compositionality of microbiome data and minimize the occurrence of false positives through the careful application of multiple test corrections (20, 49).

PERMANOVA statistical tests were used to address whether ANXD and MDD were significantly associated with microbiota beta diversity. When both medicated and unmedicated participants were considered, MDD but not ANXD, was significantly associated with microbiota beta diversity but when only unmedicated participants were considered, there was no significant association between either MDD or ANXD and microbiota beta diversity. This suggests that some of the differences in the microbiota between participants with MDD and HC was due to medication-induced microbiota differences. This finding is consistent with some previous studies; when only unmedicated participants were included, there is no significant difference but when medication use is considered, there is a significant difference (21, 61).

We identified biomarkers of ANXD, MDD, anxiolytic use, and antidepressant use and validated them using log ratios. Interestingly, there was a large overlap between microbes associated with ANXD and those associated with MDD. There were 31 microbes credibly associated with both ANXD and MDD, and 31 different microbes credibly associated with HC relative to both ANXD and MDD (**Fig S1A-B**). The log ratio constructed from these common microbes was significantly enriched in participants with ANXD, MDD, and both diagnoses relative to HC. In participants with both diagnoses relative to MDD only, the enrichment held regardless of medication use (**Fig S1E-F**). However, there was only a small overlap between ASVs associated with anxiolytics and antidepressants use (**Fig S1C-D**). Despite this small overlap, the log ratio credibly differentiated subjects using both anxiolytics and antidepressants,with unmedicated participants. It further differentiated those using both anxiolytics and antidepressants, for those only using antidepressants. (**Fig S1G**).

There were minimal overlaps between microbes associated with antidepressants or anxiolytics use and those associated with ANXD or MDD (**Fig S2**). One notable exception is there were twelve microbes that were both significantly associated with antidepressant use and HC (relative to MDD; **Figure S2D**). This observation is somewhat consistent with previous reports that taking medications for ANXD and/or MDD returns the gut microbiota to a “normal” state (10, 11). However, given that there are several more microbes that are unique to participants using antidepressants, a more parsimonious explanation may be that antidepressant use restores levels of some microbes that are depleted due to MDD. There is still a large number of microbes that are depleted after antidepressant use, including several that are not depleted due to MDD alone. This suggests that microbes, or microbiota-altering interventions (such as dietary interventions) could be used in tandem with medication use to restore microbes depleted by both MDD itself and antidepressants.

This study notably also identifies microbes characteristic of ANXD that can distinguish not only between participants with ANXD and HC but also participants with both ANXD and MDD and participants with MDD alone (**Fig 2C**). This is significant as there has been considerably less research on the GMBA in ANXD compared to the GMBA in MDD, bipolar disorder, and schizophrenia. However, the ANXD analysis is constrained by the fact that the American Gut cohort cannot be used as a validation cohort because no information was collected on ANXD diagnoses. Further primary literature on changes in the gut microbiome in response to ANXD is needed.

The interplay between ANXD, MDD, lifestyle factors, and the microbiota highlights a complex network of interactions that underscore the multifaceted nature of NPDs. ANXD and MDD often coexist, suggesting a shared pathophysiological pathway that may be influenced by the gut microbiota. Lifestyle factors such as diet, exercise, stress levels, and sleep patterns further modulate this relationship, potentially altering the gut microbial composition and its metabolic outputs, which in turn can affect brain function and emotional regulation through the gut-brain axis. For instance, diets rich in prebiotics and probiotics can favorably modify the gut microbiota, potentially reducing inflammation and altering neurotransmitter activity, thereby mitigating symptoms of anxiety and depression. Similarly, regular physical activity has been shown to influence microbial diversity, enhancing the abundance of beneficial microbes that produce mood-regulating compounds like short-chain fatty acids. This dynamic interplay suggests that interventions targeting lifestyle modifications, alongside strategies for modulating the gut microbiome, could offer a comprehensive approach to managing ANXD and MDD, emphasizing the need for integrated treatment strategies that consider both the microbiome and lifestyle factors in their therapeutic protocols.

The evidence of a relationship between the gut microbiota and NPD, including the nuanced effects of psychiatric medications on microbial diversity, opens new avenues for clinical practice. Recognizing specific microbial markers associated with ANXD and MDD paves the way for the development of novel diagnostic tools and therapeutic strategies, potentially allowing for more personalized and effective treatment plans. The modulation of the gut microbiome through targeted probiotic interventions or dietary modifications could complement or even reduce reliance on traditional pharmacotherapy, mitigating side effects and improving overall treatment outcomes. Furthermore, understanding the role of microbiome offers a compelling rationale for integrating gut health assessments into routine psychiatric evaluations, encouraging a holistic approach to mental health care. This paradigm shift towards incorporating microbiome analysis in clinical settings could revolutionize the management of NPD, emphasizing the importance of gut health in mental well-being and opening up a new frontier in psychiatric treatment and prevention strategies.

This study’s strengths include employing compositionally aware methods on a large, United States-based cohort; however, it is not without its limitations. The study does not account for medication dosage variations, and while it distinguishes between anxiolytics and antidepressants use, it does not differentiate between different drugs or dosages within these medication categories to maintain statistical power. The reliance on self-reported medication usage introduces potential inaccuracies regarding adherence to prescribed regimens or unsupervised medication consumption. Another limitation arises from the methodological choice of utilizing 16S rRNA gene amplicon data over shotgun metagenomic sequencing. While shotgun metagenomic data was available for this study and microbial signatures in this data were able to consistently differentiate between participants with ANXD and MDD, and HC, other subgroups were harder to distinguish (**Fig S3**). This is likely due to the limited sample size, as we saw smaller differences in the same 16S data subsetted to participants with both a 16S sample and a WGS sample (**Fig S3**). This suggests a need for future research to prioritize shotgun metagenomic data, which is becoming more cost-effective and informative. Finally, the machine learning techniques employed in this study, while they effectively distinguish between HC and participants with ANXD or MDD, they do not represent the cutting edge in data analysis. Advanced deep learning models could potentially offer finer distinctions among participant groups. However, such models often lack transparent feature selection, which could impede their practical applicability in clinical settings due to challenges in interpreting which specific features drive observed differences. Future research might benefit from integrating untargeted approaches, such as metabolomics, to classify participants more accurately by medication use, enhancing the study’s depth and the reliability of its conclusions.

## 8. Acknowledgements

We thank Caitriona Brennan, MacKenzie Bryant, Lindsay DeRight Goldasich, Karenina Sanders, and Julia Toronczak for 16S preparation.

The Tulsa study is led by Dr. Martin Paulus. We thank the participants of the Tulsa-1000 cohort and the NeuroMAP CoBRE study for their samples.

This work was carried out by the Alzheimer Gut Microbiome Project (AGMP) an initiative funded by National Institutes on Aging (U19AG063744) led by Rima Kaddurah Daouk, Rob Knight, and Sarkis Mazmanian in partnership with large number of academic institutions. The investigators within the AGMP not listed specifically in this publication’s author’s list provided insights that helped with study design, analysis of data, dealing with confounding factors among other contributions that enabled analysis or writing of this manuscript. A listing of AGMP Investigators can be found at https://alzheimergut.org/meet-the-team/. All data from the U19 initiative is or will be posted in AD Knowledge Portal (Sage Bionetworks). The work was funded wholly or in part by the following grants and supplements thereto awarded to Rima Kaddurah Daouk : NIA R01AG046171, RF1AG051550, RF1AG057452, R01AG059093, RF1AG058942, U01AG061359.

---

**Figure S1.**
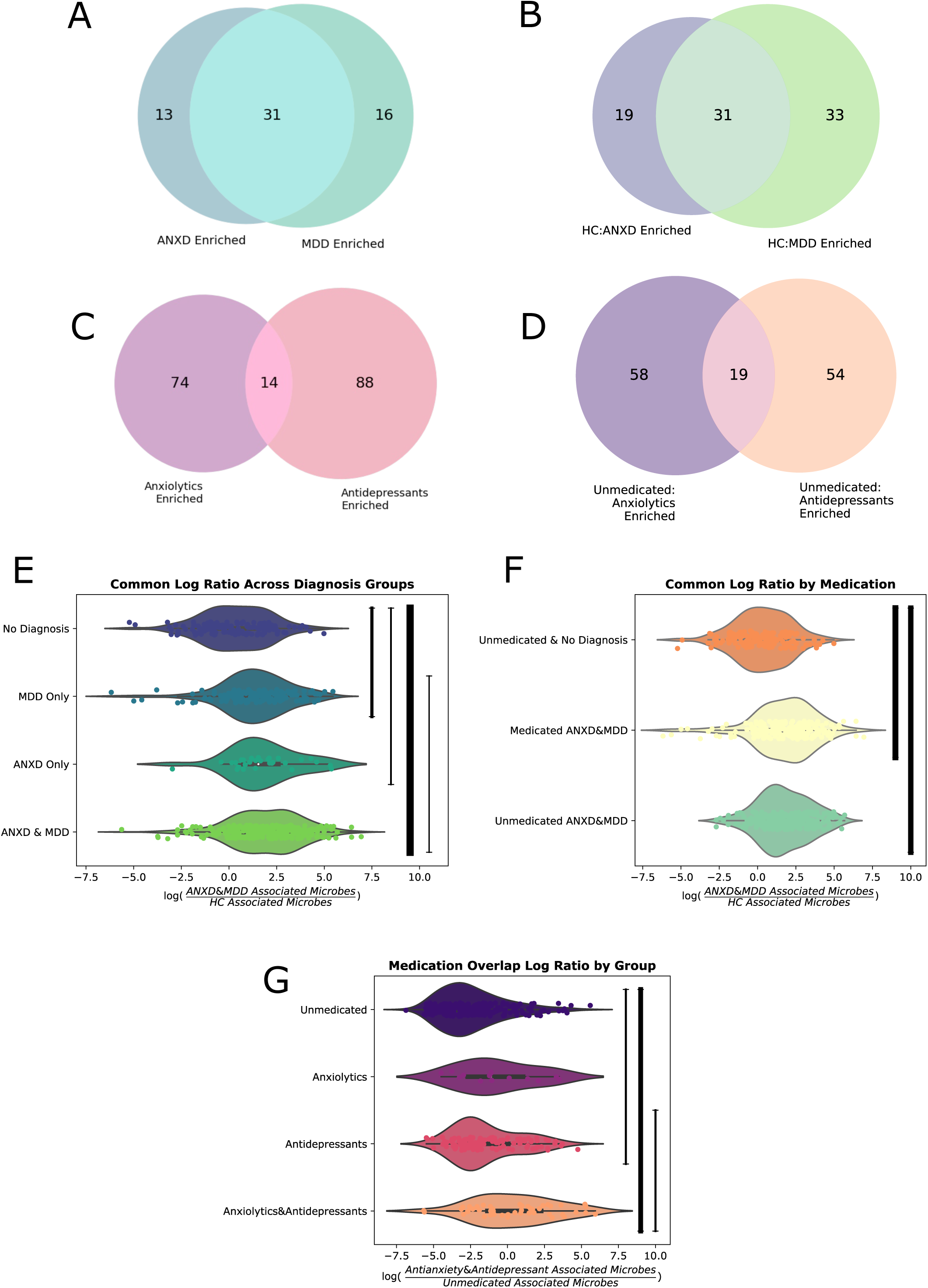
ANXD and MDD have similar effects on gut microbiota; anxiolytics and antidepressants have more distinct effects. **(A)** Venn diagram showing overlap in the credible microbes associated with ANXD and MDD. **(B)** Venn diagram showing overlap in the credible microbes associated with HC relative to ANXD and HC relative to MDD. **(C)** Venn diagram showing overlap in the credible microbes associated with anxiolytics and antidepressants. **(D)** Venn diagram showing overlap in the credible microbes associated with unmedicated participants relative to those on anxiolytics and unmedicated participants relative to those on antidepressants. **(E)** A log ratio was created using the microbes that were credibly associated with both ANXD and MDD and the microbes associated with HC relative to both ANXD and MDD. This log ratio was elevated in participants with MDD only, ANXD only, and ANXD and MDD relative to HC. It was also enriched in participants with ANXD and MDD relative to MDD alone. **(F)** This “common” credible log ratio was also enriched in both unmedicated and medicated participants with ANXD and MDD relative to unmedicated participants. **(G)** Another log ratio was created using the microbes that were credibly associated with both anxiolytics and antidepressants and the microbes associated with unmedicated participants relative to both anxiolytics and antidepressants.

**Figure S2.**
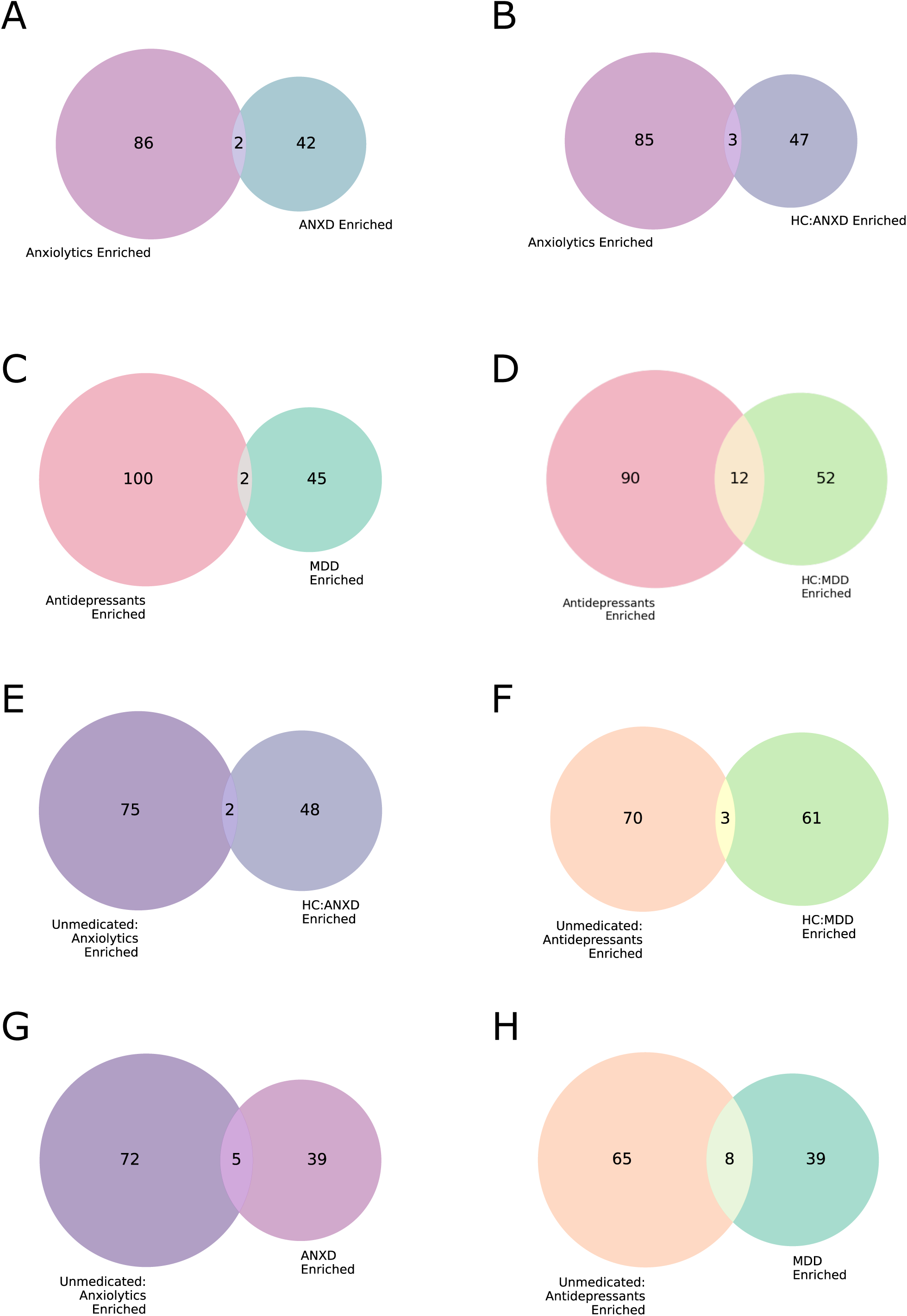
Little overlap in microbes enriched in medications and those enriched in psychiatric disease. **(A)** Venn diagram showing minimal overlap in the credible microbes associated with ANXD and anxiolytics. **(B)** Venn diagram showing minimal overlap in the credible microbes associated with HC relative to ANXD and anxiolytics. **(C)** Venn diagram showing minimal overlap in the credible microbes associated with MDD and antidepressants. Venn diagram showing some overlap in the credible microbes associated with HC relative to MDD and antidepressants. **(E)** Venn diagram showing minimal overlap in the credible microbes associated with HC relative to ANXD and unmedicated participants relative to those on anxiolytics. **(F)** Venn diagram showing minimal overlap in the credible microbes associated with HC relative to MDD and unmedicated participants relative to those on antidepressants. **(G)** Venn diagram showing minimal overlap in the credible microbes associated with ANXD and unmedicated participants relative to those on anxiolytics. **(H)** Venn diagram showing some overlap in the credible microbes associated with MDD and unmedicated participants relative to those on antidepressants.

**Figure S3.**
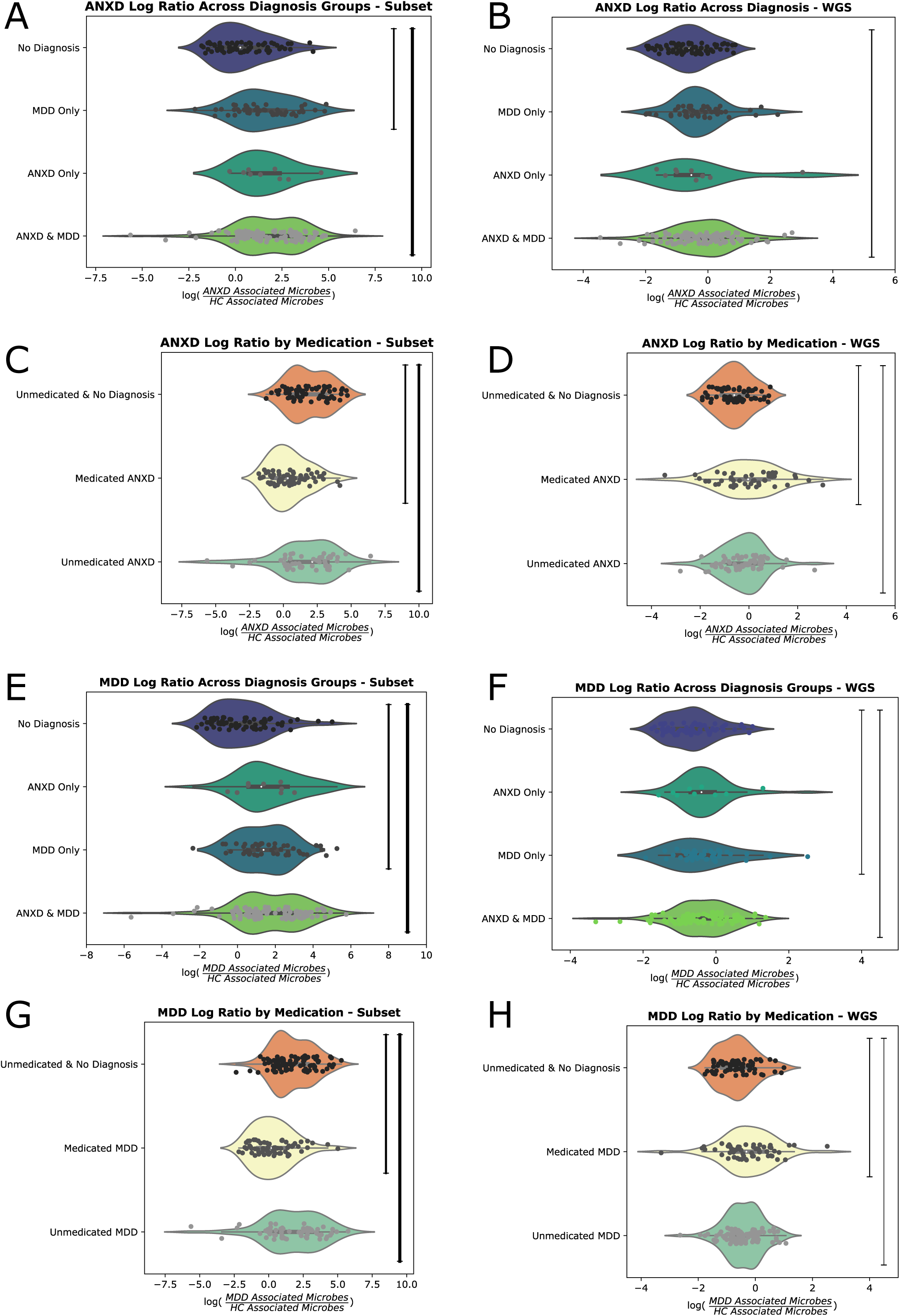
Smaller differences in enriched microbes in metagenomic data may be explained by the differences in the subset themselves. **(A)** The log ratio of microbes credibly associated with ANXD relative to microbes credibly associated with HC displayed only in participants who have a matching sample sequenced by WGS as well. In this subset, there is a significant difference in participants with MDD only and participants with ANXD and MDD relative to HC. **(B)** We used nearest neighbors on the Greengenes2 (GG2) phylogenetic tree to transfer our credibly associated microbes with 16S data to WGS data. This WGS log ratio for ANXD was enriched only in participants with both ANXD and MDD. **(C)** The ANXD-specific log ratio was significantly lower in medicated participants with ANXD and significantly higher in unmedicated participants with ANXD relative to unmedicated HC. **(D)** The WGS ANXD-specific log ratio was significantly higher in both medicated and unmedicated participants with ANXD relative to unmedicated HC. **(E)** The log ratio of microbes credibly associated with MDD relative to microbes credibly associated with HC displayed only in participants who have a matching sample sequenced by WGS as well. In this subset, there is a significant difference in participants with MDD only and participants with ANXD and MDD relative to HC. **(F)** We used nearest neighbors on the GG2 phylogenetic tree to transfer our credibly associated microbes with 16S data to WGS data. This WGS log ratio for MDD was enriched in participants with MDD alone and those with ANXD and MDD relative to HC. **(G)** The MDD-specific log ratio was significantly lower in medicated participants with MDD and significantly higher in unmedicated participants with MDD relative to unmedicated HC. **(H)** The WGS MDD-specific log ratio was significantly higher in both medicated and unmedicated participants with MDD relative to unmedicated HC.

**Figure S4.**
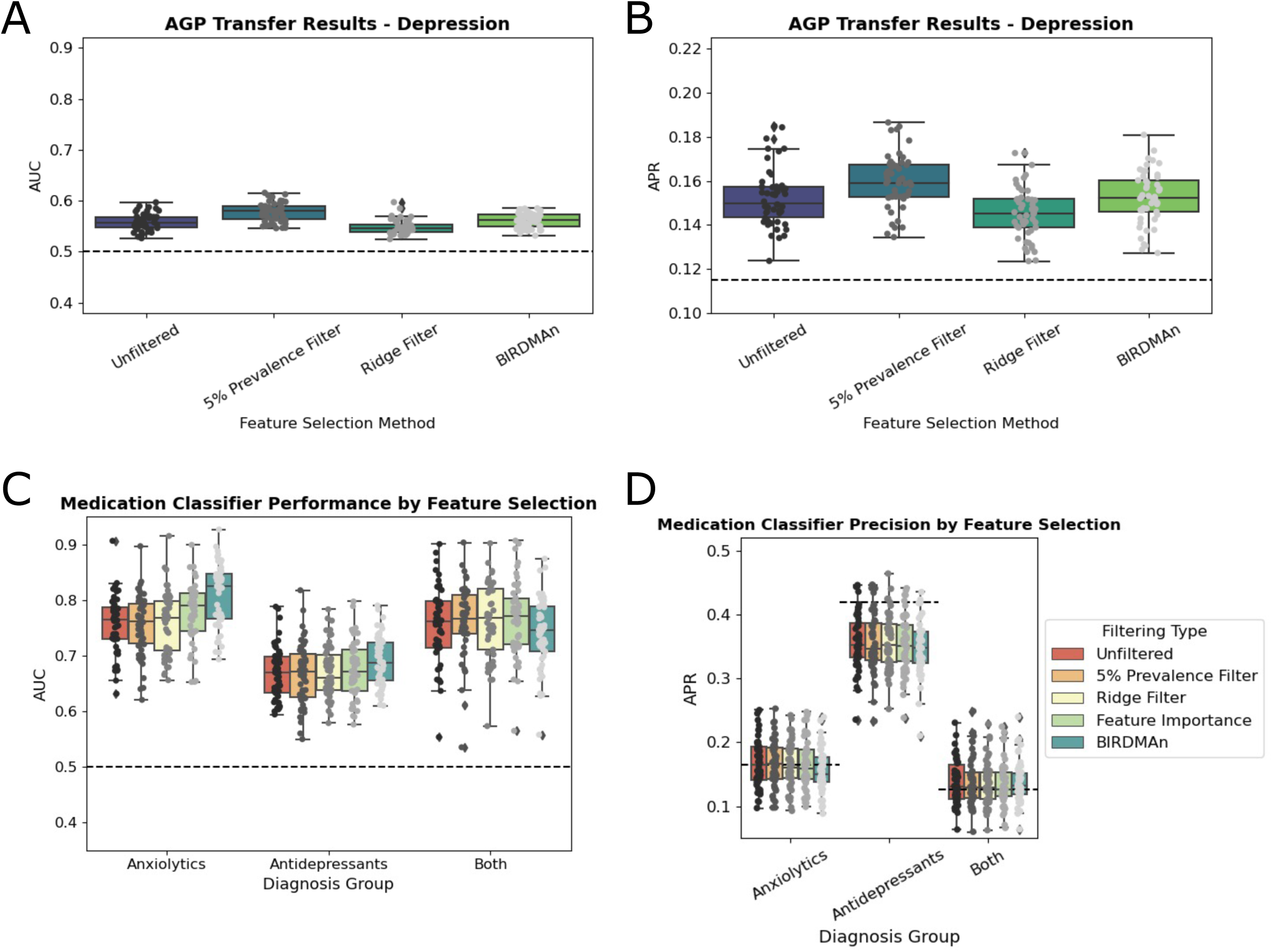
Random Forest Classifiers demonstrate high accuracy but low precision when applied to American Gut cohort and medication use. We used a Random Forest Classifier to distinguish AG participants who self-reported a MDD diagnosis from those who self-reported no diagnosis. **(A)** Area under the receiver operating characteristic curve (AUC) for the AGP diagnosis classifier. **(B)** Average precision scores (APR) for the AGP diagnosis classifier. **(C)** AUCs for a medication classifier across the cohort. **(D)** APRs for the medication classifier.

